# The Neuro-Immune Axis in Glioma Spontaneous Remission: Evidence from Population-Level Latent Variable Modelling and Transcriptomic Profiling

**DOI:** 10.64898/2026.06.19.733335

**Authors:** Ananta Kapoor, Anil Kumar Tiwari, Siddharth Srivastava

## Abstract

Gliomas are primary brain tumours that develop from neural stem or progenitor cells containing oncogenic alterations. Gliomas undergo remission with partial or complete disappearance of the disease, and in rare instances, spontaneous remission. Spontaneous remission happens without treatment or with inadequate medical intervention. This is a rare but well-documented phenomenon and has been observed across various tumour types, including gliomas, with varying frequency. While historically viewed as clinical anomalies, we hypothesize that SR may be driven by a measurable, latent neuro-immune axis specifically, autonomic vagal nerve modulation of the tumour microenvironment (TME) via the cholinergic anti-inflammatory pathway. Using a Gamma frailty Cox proportional hazards model on a SEER cohort of 6,939 glioma patients, we identified a latent biological variable (Z) that explains 27.1% of survival variance independent of age, grade, treatment, and tumour location. To determine the molecular basis of this latent survival advantage, we applied parallel frailty and Cox models to the TCGA Lower Grade Glioma and Glioblastoma (LGG+GBM) cohorts. Clinical validation confirmed expected hazards for age and grade, with the frailty model achieving high predictive accuracy (5-year AUC = 0.840; 10-year AUC = 0.841). Transcriptomic integration revealed that the alpha7 nicotinic acetylcholine receptor (CHRNA7) is highly protective, inversely correlating with biological frailty (r = -0.285, p < 0.0001). Conversely, pro-inflammatory cytokines (IL6) and M2 macrophage markers (CD163) positively correlated with frailty. Grade-stratified Cox regression and Kaplan-Meier analyses confirmed that CHRNA7 confers a significant survival advantage entirely independent of tumour grade. Single-cell RNA-sequencing data (Core GBmap) confirmed that CHRNA7 and TLR4 are expressed heavily on tumour-associated macrophages and microglia, rather than malignant cells. Our in-silico integration suggests that high vagal tone releases acetylcholine, binding to alpha7nAChR on TME macrophages. This triggers a signalling cascade that dampens the IL-6 production required for glioma proliferation, effectively halting tumour growth. Ongoing in vitro wet-lab experiments utilizing specific alpha7nAChR agonists (GTS-21) and physiological stress models aim to clinically validate this vagal-immune mechanism.

## Introduction

Gliomas are the most common primary brain tumours, characterized by aggressive infiltration and dismal prognoses. Despite advances in surgery, radiation, and chemotherapy, survival rates remain low. However, in exceptionally rare instances, gliomas undergo spontaneous remission (SR) the partial or complete disappearance of a tumour in the absence of targeted or adequate treatment. These cases are often dismissed as clinical anomalies or misdiagnoses. We hypothesize that SR is instead the extreme manifestation of an endogenous, latent biological mechanism that actively suppresses tumour growth.

Recent advances in neuro-immunology highlight the cholinergic anti-inflammatory pathway (CAP), a mechanism by which the autonomic nervous system regulates systemic inflammation. The efferent vagus nerve releases acetylcholine (ACh), which binds to alpha7 nicotinic acetylcholine receptors (alpha7nAChR) on macrophages. Activation of alpha7nAChR is an essential regulator of inflammation, triggering the JAK2-STAT3 signalling cascade to potently inhibit the release of pro-inflammatory cytokines such as IL-6 and TNF-alpha. Furthermore, pathological stress states and damage-associated molecular patterns (DAMPs) upregulate alpha7nAChR on macrophages, priming them for cholinergic regulation.

In the glioma tumour microenvironment (TME), the immune system is actively hijacked. Tumours exploit chronic inflammation particularly via IL-6 to drive proliferation and invasion, while simultaneously promoting an M2-polarized macrophage state (clinically marked by CD163) to evade clearance. We propose that high vagal tone, which is clinically measurable via Heart Rate Variability (HRV) and acts as an independent prognosticator for prolonged cancer survival, serves as a systemic disruptor of this pro-tumour TME. By maintaining high cholinergic signalling, the vagus nerve engages *α*7nAChR on TAMs. Based on established neuro-immune mechanisms, this receptor activation specifically suppresses the release of pro-tumorigenic cytokines like IL-6. We hypothesize that this targeted cytokine suppression deprives the glioma of critical inflammatory growth factors, effectively altering the ‘soil’ of the TME. In extreme cases, this neuro-immune modulation may so profoundly restrict tumor progression that it halts growth entirely, manifesting clinically as spontaneous remission. In this study, we utilize large-scale epidemiological modelling (SEER database) alongside bulk (TCGA) and single-cell (CellxGene) transcriptomics to map this neuro-immune axis in glioma, seeking to decode the mechanisms of vagus nerve influencing macrophages through the cholinergic anti-inflammatory pathway resulting in a better prognosis.

## Methods

### SEER Cohort Frailty Modelling

Clinical and survival data for 6,939 glioma patients were obtained from the Surveillance, Epidemiology, and End Results (SEER) database. A Gamma frailty Cox proportional hazards model was applied to adjust for age, sex, grade, surgery, chemotherapy, radiation, and tumour location. The frailty weight (Z) was extracted to quantify latent biological vigour exceeding clinical predictions.

### TCGA Clinical-Genomic Frailty and Survival Models

Bulk RNA-seq data and clinical metadata for LGG and GBM cohorts were accessed via TCGA. A multivariable Cox proportional hazards model was constructed using age, sex, grade, histology, radiation, and IDH1 mutation status. Frailty Z scores were extracted for each patient. We calculated Spearman rank correlations between Frailty Z and the expression of neuro-immune genes (CHRNA7, IL6, IL10, CD163, S100B, TLR4, P2RY12, ITGAM). To ensure transcriptomic signals were not confounded by tumour grade, we generated grade-stratified Cox regression models and compared gene expression between top 25 percent and bottom 25 percent survival cohorts within each grade.

### Single-Cell RNA-Sequencing (scRNA-seq)

To resolve the cellular origin of our target genes within the TME, we utilized the Harmonized single-cell landscape of glioblastoma (Core GBmap) via the CZ CELLxGENE platform. Uniform Manifold Approximation and Projection (UMAP) and violin plots were generated to assess the expression of CHRNA7, TLR4, TMEM119, and ITGAM across malignant cells, macrophages, microglia, and astrocytes.

## Results

### A Latent Biological Variable Explains Spontaneous Remission Independent of Tumour Location

Using the SEER cohort (n = 6,939), we investigated the existence of an unmeasured biological variable driving survival. Our Gamma frailty Cox model revealed a residual unexplained variance of 0.271, indicating that 27.1 percent of survival variance remains unexplained by standard clinical metrics. Notably, SR cases (n = 14) achieved 100 percent survival over 266 to 662 months, vastly outperforming the SEER median. These SR cases clustered tightly around a frailty value of Z = 1 even after adjusting for 12 distinct tumour locations. Furthermore, within-location analysis of optic nerve tumours showed that while non-SR patients experienced standard mortality, SR cases at the exact same anatomical site exhibited zero mortality and significantly lower frailty (p < 0.00001). This proves that SR is driven by a systemic biological factor, not localized anatomical advantages.

**Figure 1.**
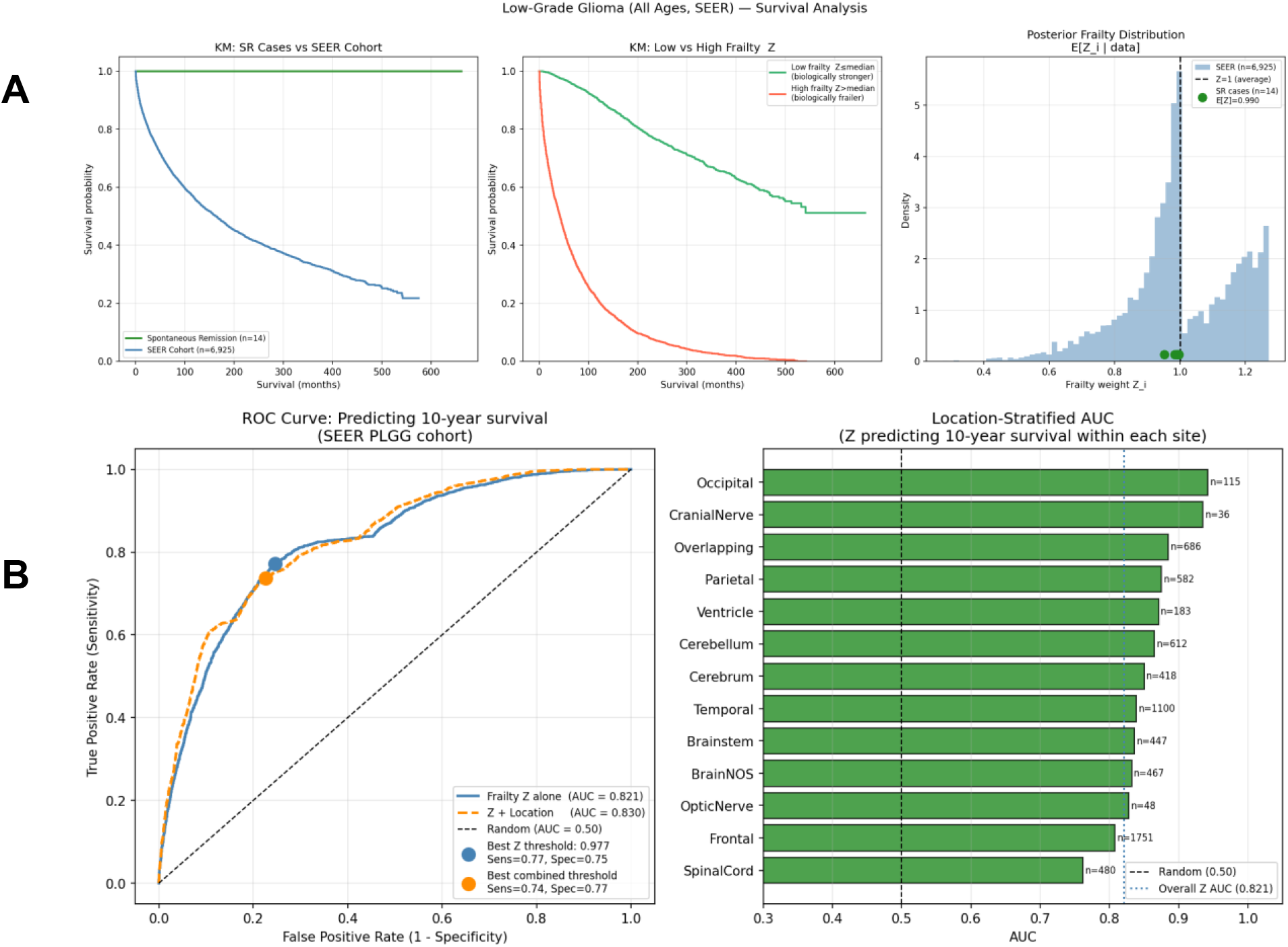
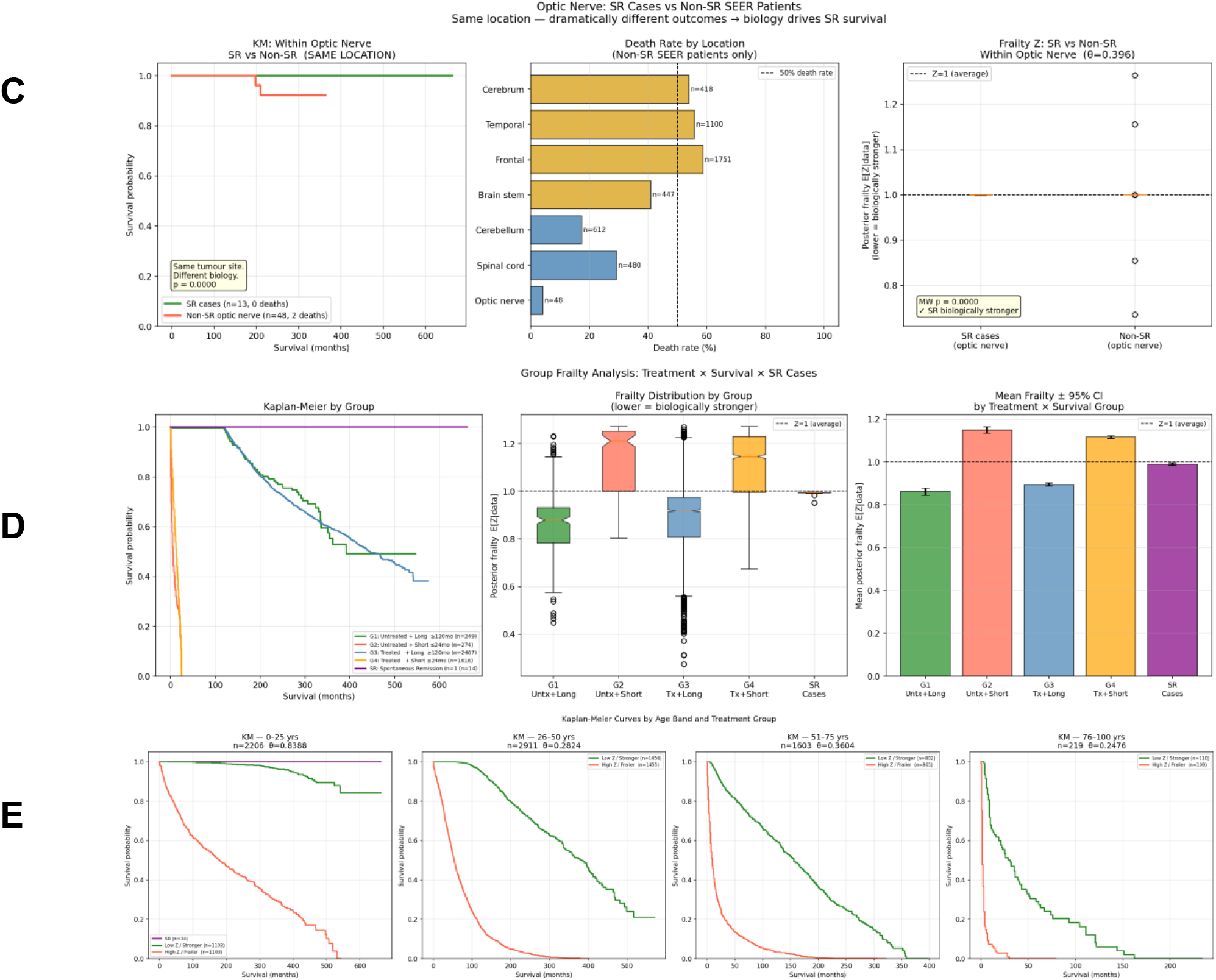
Latent biological frailty in the SEER glioma cohort. **(A)** Gamma frailty Cox proportional hazards model demonstrating survival distribution across treatment groups. **(B)** ROC curve analysis showing high predictive accuracy (AUC > 0.82) of the frailty metric for 10-year survival. **(C)** Within-location analysis of optic nerve tumors, proving spontaneous remission cases possess distinct biological vigor independent of tumor anatomy. **(D)** Group frailty analysis with treatment **(E)** Age-stratified frailty analysis confirming the survival advantage persists independently across all age bands.

### Clinical Validation of the Frailty Model in the TCGA Cohort

To uncover the molecular identity of this latent biological variable, we applied our Frailty Cox regression to the TCGA dataset. The model successfully replicated SEER findings, demonstrating robust discrimination for long-term survival with a 5-year AUC of 0.840 and a 10-year AUC of 0.841. The model accurately captured known clinical hazards: older age and higher grade significantly increased hazard risk, while Oligodendroglioma histology and IDH1 mutations were highly protective. Age-stratified and group-frailty analyses confirmed that the latent Frailty Z score effectively distinguishes long-survivors from short-survivors across all age bands and histological subtypes.

**Figure 2.**
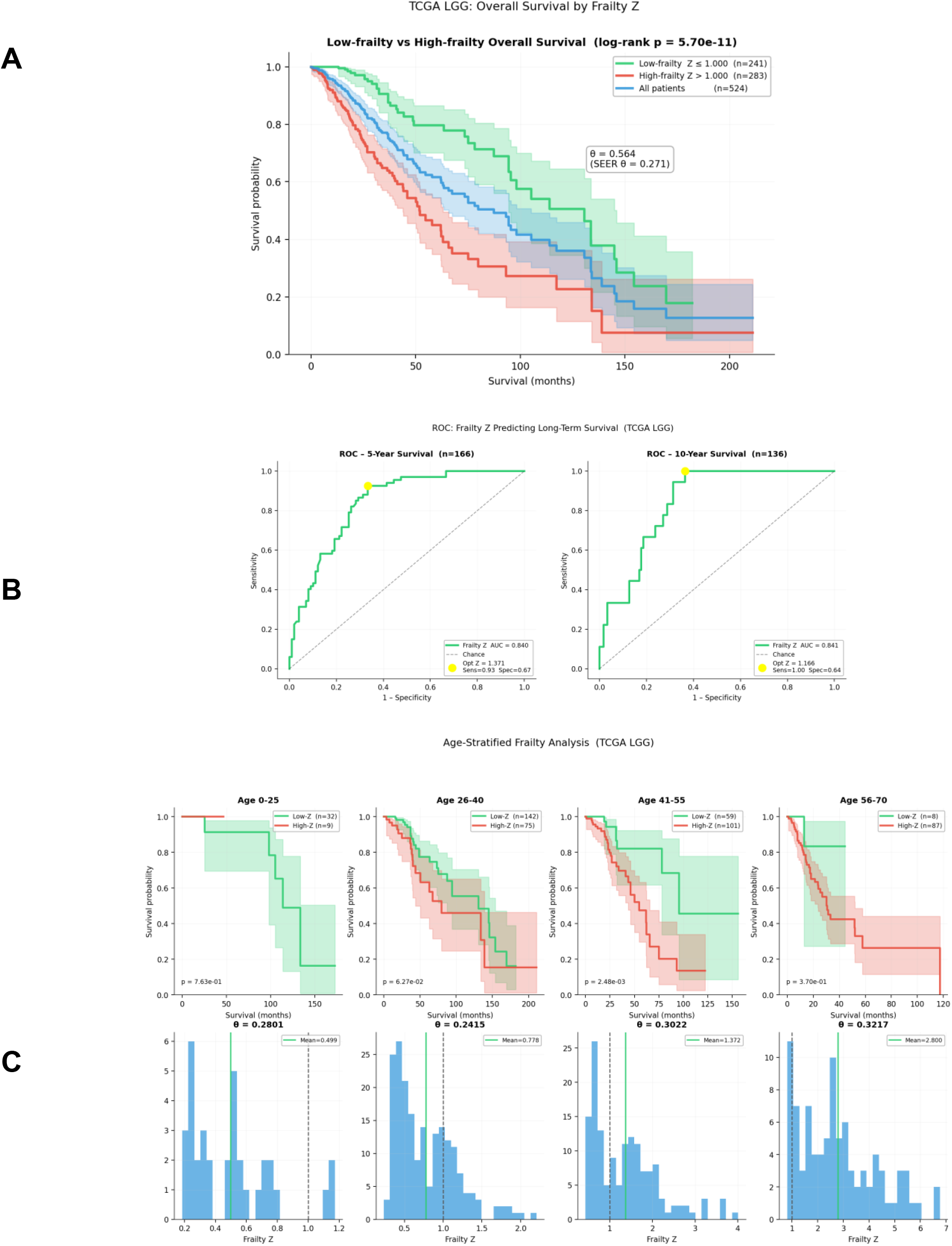
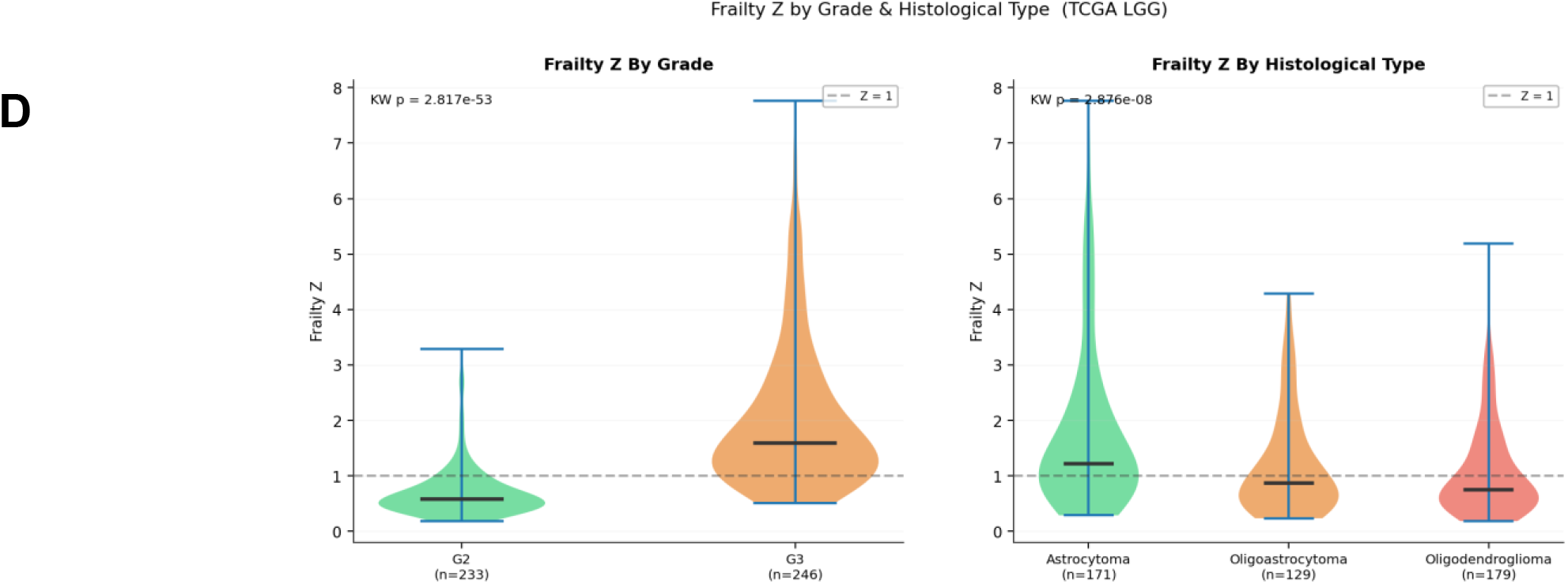
Clinical validation of the frailty model in the TCGA cohort. **(A)** Kaplan-Meier overall survival curve stratifying TCGA LGG patients by High versus Low Frailty Z scores. **(B)** Receiver Operating Characteristic (ROC) curves confirming the model’s accuracy for 5-year and 10-year survival. **(C)** Age-stratified Frailty Z distribution, demonstrating effective separation of survivors across distinct age cohorts. **(D)** Violin plots displaying Frailty Z score distribution across WHO grades and specific histological subtypes. **(E)** Kaplan-Meier overall survival curve stratifying patients by IDH1 mutation status.

### Transcriptomic Profiling Identifies CHRNA7 as a Potent Marker of Biological Vigor

When correlating patient Frailty Z scores with gene expression, the cholinergic alpha7nAChR receptor (CHRNA7) emerged as the most significant protective factor. CHRNA7 expression was strongly inversely correlated with frailty (r = -0.285, p = 1.96e-10), indicating that patients with higher expression of this vagal receptor possess significantly greater biological vigour. Conversely, the pro-inflammatory cytokine IL6 (r = 0.165, p = 0.0002), the M2 tumour-associated macrophage marker CD163 (r = 0.182, p < 0.0001), and the damage-associated molecular pattern S100B (r = 0.190, p < 0.0001) all showed significant positive correlations with frailty, marking them as drivers of tumour lethality. A correlation heatmap of these targets confirmed a tightly co-regulated network linking the cholinergic and inflammatory systems within the tumour microenvironment.

**Figure 3.**
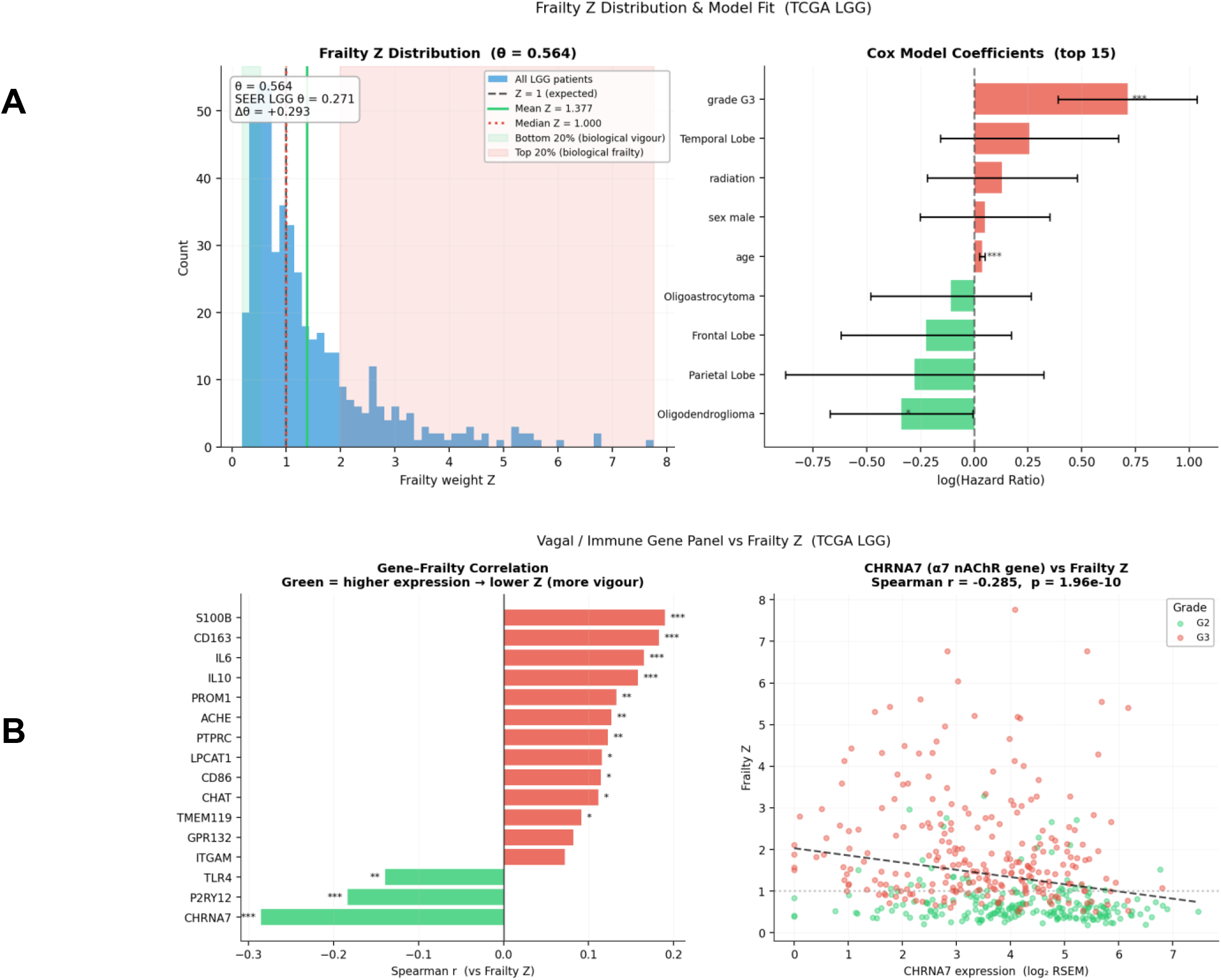
Transcriptomic correlation with biological frailty. **(A)** Multivariable Cox model coefficients confirming standard clinical hazards (age, grade, radiation) alongside the Frailty Z metric. **(B)** Spearman rank correlation bar chart identifying CHRNA7 as the most protective gene (negative correlation) and IL6/CD163 as harmful drivers (positive correlation).

### The Cholinergic Survival Advantage is Independent of Tumour Grade

Because lower-grade tumours inherently survive longer and might express different baseline genes, we sought to prove that CHRNA7’s protective effect is independent of tumour grade. A grade-stratified Cox regression model, alongside combined clinical-genomic forest plots, confirmed that CHRNA7 exerts a protective hazard ratio even when grade is strictly controlled. Kaplan-Meier analysis further demonstrated that high CHRNA7 expression significantly prolongs survival overall (p = 0.0023) and maintains protective trends within specific tumour grades. Furthermore, boxplot analysis of top 25 percent (long) versus bottom 25 percent (short) survivors revealed that CHRNA7 is significantly upregulated in the longest survivors, specifically within Grade 3 cohorts (p = 0.047).

**Figure 4.**
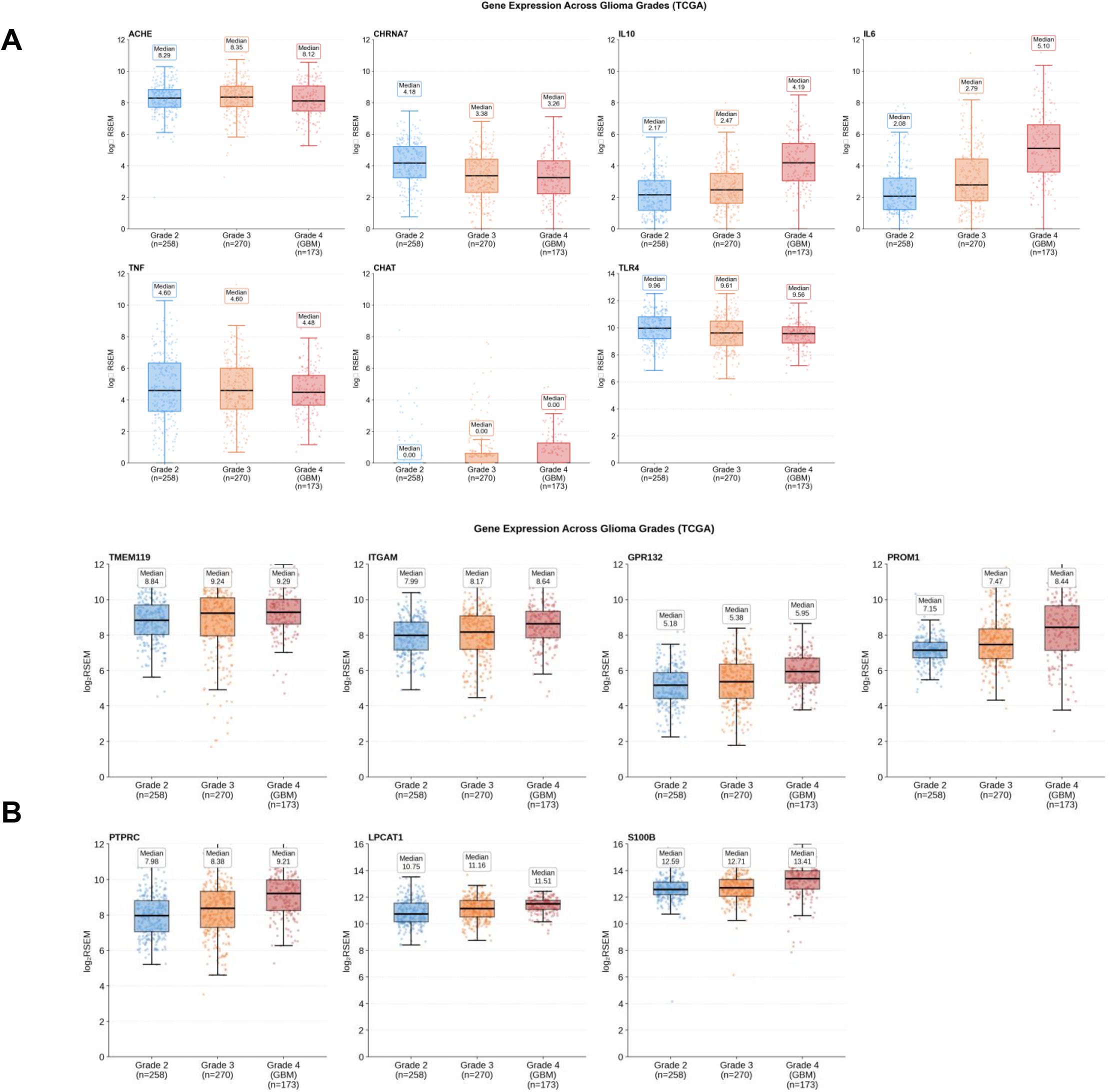
Expression of neuro-immune and inflammatory genes across glioma progression. (A) Gene expression profiles (log_2_RSEM) of cholinergic pathway components (*CHRNA7, ACHE, CHAT*) and key inflammatory regulators (*IL6, TNF, IL10, TLR4*) across WHO Grades 2, 3, and 4 (GBM). (B) Expression distribution of resident microglial (*TMEM119*), macrophage (*ITGAM*), and astrocyte/stress (*S100B*) markers across tumor grades, demonstrating the evolving immune landscape of the tumor microenvironment.Single-Cell Resolution Identifies Macrophages and Microglia as Targets of Cholinergic Regulation

To confirm that these cholinergic receptors are present on immune cells rather than the tumour itself, we queried the GBmap single-cell dataset. Multi-metric profiling and UMAP clustering revealed that CHRNA7 expression is localized strictly to specific subsets of the myeloid compartment (macrophages and microglia) and neurons, rather than malignant glioma cells. Similarly, TLR4 and ITGAM were heavily clustered in the tumour-associated immune cell populations. This confirms that the cellular infrastructure required for vagus-nerve-mediated immune suppression exists robustly within the glioma TME.

**Figure 5.**
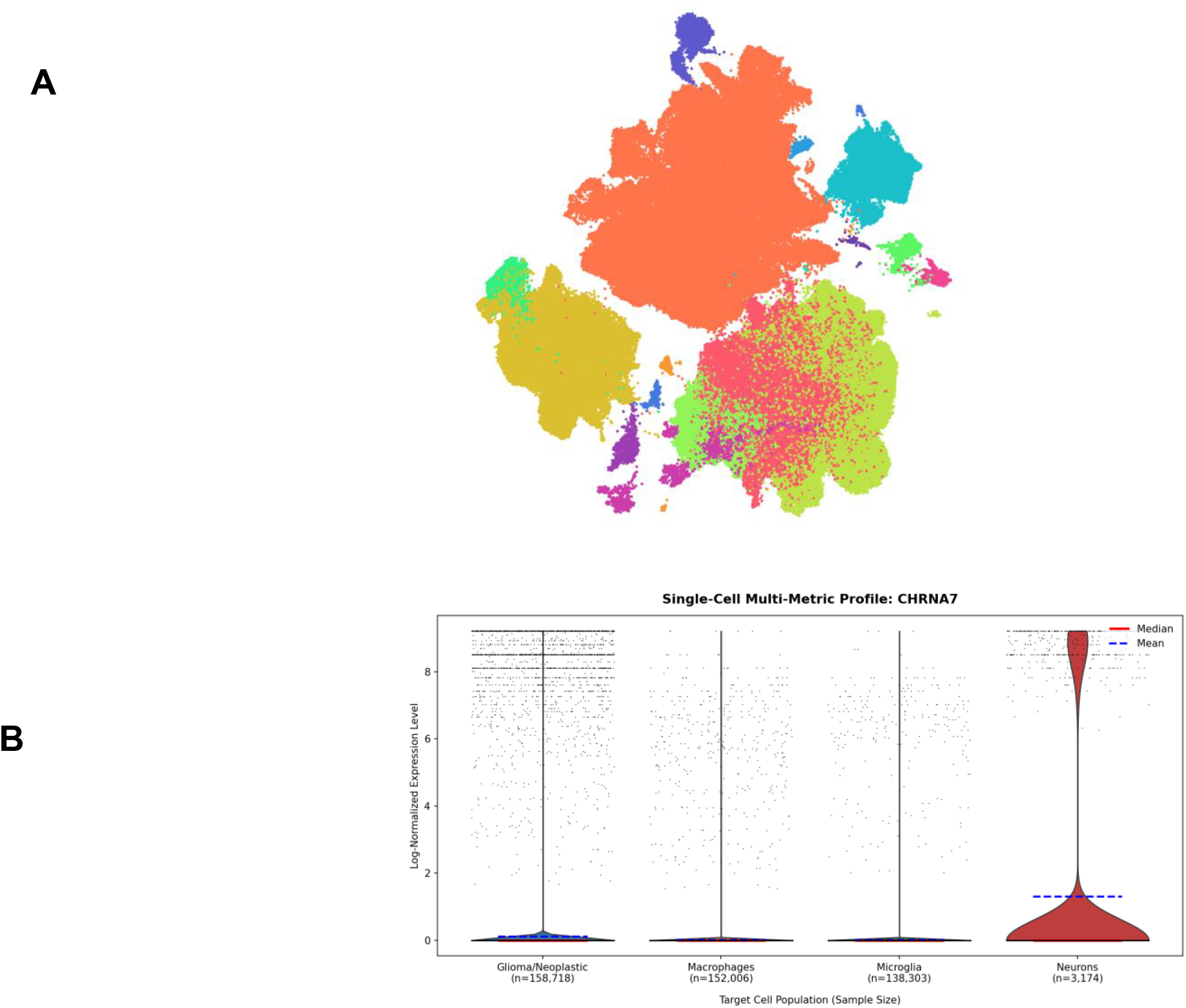
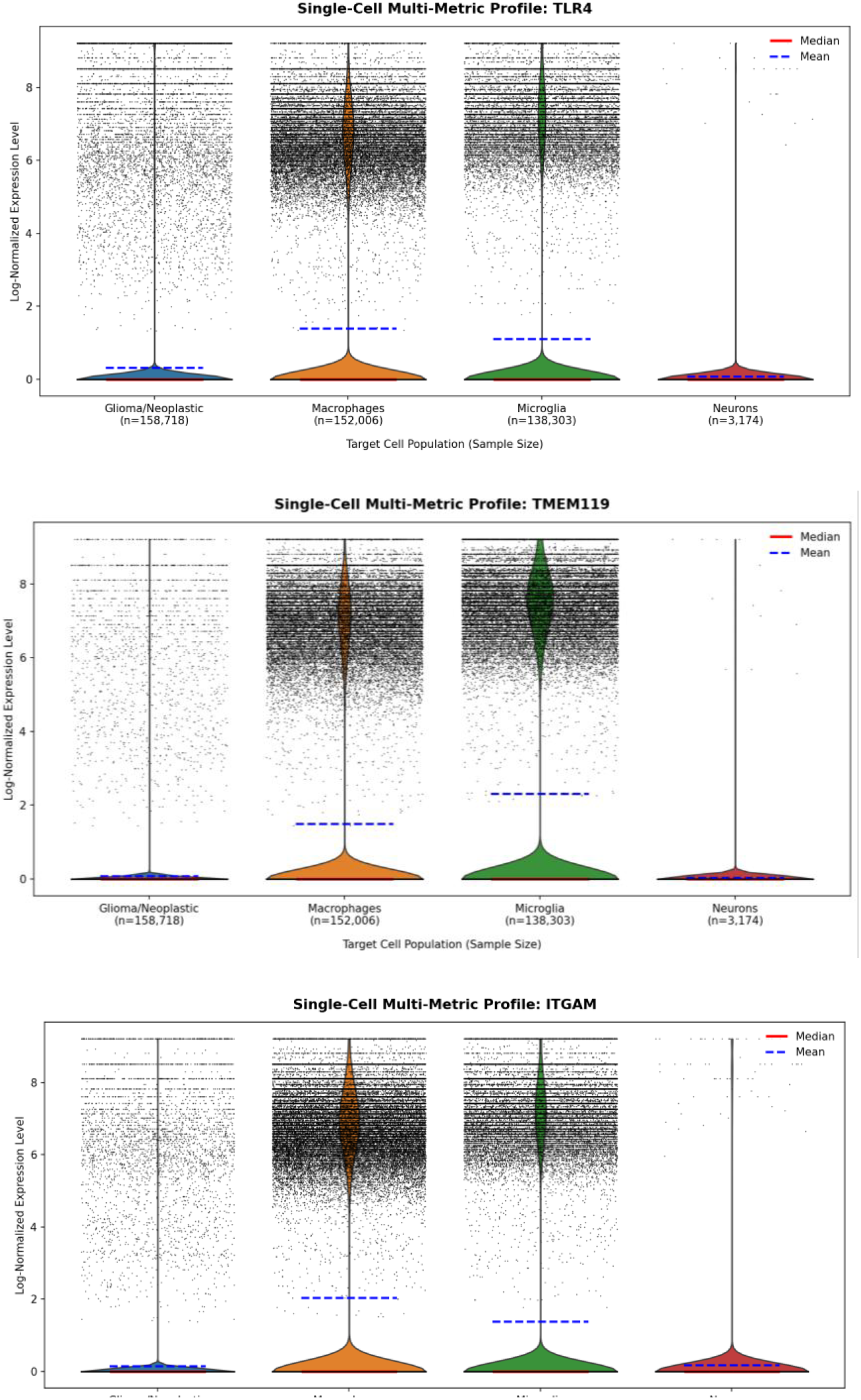
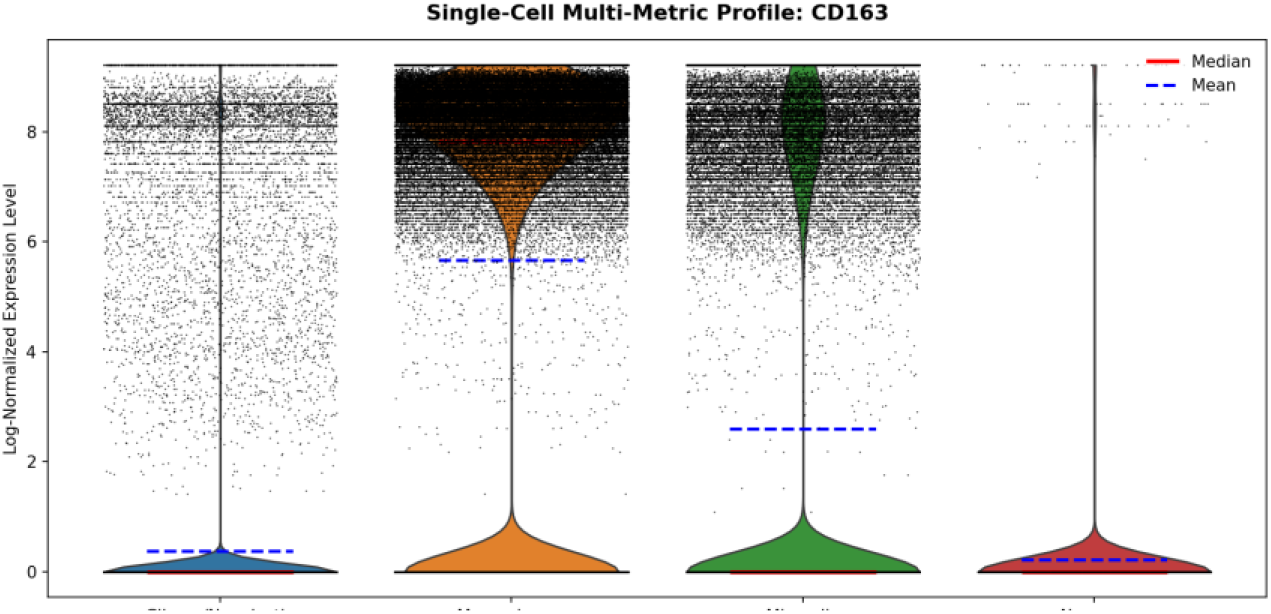
Single-cell resolution maps cholinergic targets to TME macrophages and microglia. (A) Global UMAP projection of the Harmonized Core GBmap dataset representing 338,564 individual cells. Each dot represents a single cell, clustered by transcriptomic similarity. Colors indicate distinct cell lineages: malignant glioma cells (orange), macrophages (red), microglia (yellow-green), mature T cells (yellow), and oligodendrocytes (cyan). (B) Multi-metric violin plots confirming *CHRNA7, TLR4, TMEM119, IITGAM, CD163* expression is significantly enriched in macrophages and microglia compared to neoplastic cells.

## Discussions and Conclusions

Integrating macroscopic epidemiological data with microscopic transcriptomic and single-cell profiling reveals a highly cohesive narrative regarding tumour suppression.

At the population level, our SEER analysis proves that over 27 percent of a glioma patient’s survival is dictated by a latent, systemic variable entirely independent of modern clinical interventions. When translated to the transcriptomic level in TCGA, this latent survival advantage neatly aligns with the cholinergic anti-inflammatory pathway. The profound inverse correlation between CHRNA7 expression and biological frailty, coupled with grade-stratified Cox models, isolates the alpha7nAChR receptor as a central node of tumour suppression.

Simultaneously, the strong positive correlation of IL6 and CD163 with frailty confirms that glioma progression relies on a pro-inflammatory, macrophage-dominant tumour microenvironment. Our single-cell data acts as the bridge between these findings: it proves that the CHRNA7 receptors are physically located on the very macrophages driving the disease.

Therefore, we conclude that the latent variable driving Spontaneous Remission is efferent vagal tone, a metric systematically linked to cancer survival outcomes. In patients with high vagal tone, the sustained release of acetylcholine into the TME binds to alpha7nAChR on tumour-associated macrophages activating the JAK STAT signalling pathway. A strong systemic vagal tone is correlated with better prognosis and survival.

### Study Limitations

While this study utilizes highly powered, multi-scale datasets to isolate the cholinergic anti-inflammatory pathway as a driver of glioma spontaneous remission, it is subject to the limitations inherent in *in silico* retrospective analyses. Bulk RNA-sequencing (TCGA) lacks spatial resolution, meaning we cannot definitively map the physical proximity of vagal efferent terminals to the *CHRNA7*-expressing macrophages. Furthermore, while statistical adjustments were made for clinical confounders, patient heart rate variability (HRV) metrics were not explicitly recorded in the SEER or TCGA databases, requiring us to rely on the Frailty Z-score as a mathematical surrogate for vagal tone. Ongoing *in vitro* and *in vivo* studies are required to prospectively validate the tumor-suppressive capabilities of *α*7nAChR agonism in the glioma TME.

## Data Availability Statement

All datasets utilized in this study are publicly available. Patient clinical and survival data were accessed via the Surveillance, Epidemiology, and End Results (SEER) Program (seer.cancer.gov). Bulk transcriptomic and clinical data for the LGG and GBM cohorts were accessed via The Cancer Genome Atlas (TCGA) through the Xena Browser (xena.ucsc.edu). Single-cell RNA-sequencing data (Core GBmap) was accessed via the Chan Zuckerberg CELLxGENE Discover platform (cellxgene.cziscience.com). Code used for the Frailty Cox proportional hazards modelling is available from the corresponding author upon reasonable request.

## Acknowledgements

We are grateful to IIT Jodhpur for its facilities and resources. I am funded by the MoE fellowship. I thank Prof. Anil Tiwari’s, Dr. Siddharth Srivastava and Prof. Sushmita Jha’s lab and students in IIT Jodhpur for their resources and mentorship.

## Author contributions

Ananta Kapoor designed and performed the experiments (data analysis, algorithms for gamma frailty cox regression model, bioinformatics analysis, designed algorithms and experiments) S.S. and A.K.T. conceptualized the study, made improvements on data analysis and designed experiments. All authors reviewed the manuscript.

